# Identifying interactions between TDP-43’s N-terminal and RNA binding domains

**DOI:** 10.1101/2024.07.31.604183

**Authors:** David D. Scott, Lipsa Jena, Samantha Perez-Miller, Vlad Kumirov, Rajesh Khanna, May Khanna

**Affiliations:** Department of Pharmacology and Therapeutics, College of Medicine, University of Florida; Department of Chemistry and Biochemistry, College of Science, University of Arizona

**Keywords:** TDP-43, HSQC-NMR, interdomain interaction, CPMG-NMR, *de novo* structure

## Abstract

TAR DNA-binding Protein 43 (TDP-43) plays a crucial role in the pathophysiology and progression of Amyotrophic Lateral Sclerosis (ALS), affecting both familial and sporadic cases. TDP-43 is an intrinsically disordered multidomain protein that consists of an N-terminal domain (NTD_1-102_), two tandem RNA recognition motifs (RRM1_102-177_ and RRM2_191-260_), and an intrinsically disordered glycine-rich C-terminal_261-414_ domain. We previously identified a chemical probe which led to allosteric alterations between the RRM and NTD of TDP-43. We attributed these changes to potential interdomain interactions between the NTD and RRM segments. In this work, we compared the 2D [^1^H,^15^N] HSQC-NMR resonances of two constructs, TDP-43_102-260_ (RRM domain alone) against TDP-43_1-260_ (NTD linked to RRM) and observed clustered shifts in the RNA binding sites of both RRM domains. To investigate why these shifts appeared in the RRM domains, in the absence of RNA, we hypothesized that the NTD domain could be stacking on the RRM domains. Thus, we modeled NTD-RRM interactions using protein-protein docking along with *de novo* sequence-to-structure predictions of TDP-43_1-260_ that propose NTD stacking onto the RRM domains. Using Carr-Purcell-Meiboom-Gill (CPMG)-relaxation dispersion NMR spectroscopy, we demonstrated evidence of an interaction between NTD_1-102_ and RRMs_102-260_. Finally, we investigated the impact of NTD on RNA binding using 2D ^15^N-HSQC-NMR and microscale thermophoresis (MST) by titration of a short UG-rich RNA sequence and observed significant changes in RNA binding between TDP-43_102-260_ and TDP-43_1-260_, further suggesting the NTD plays a role in TDP-43 RNA interactions.

---

TAR DNA-binding Protein 43 kilodaltons (TDP-43) is a ubiquitously expressed heterogeneous ribonucleoprotein (hnRNP) implicated in the pathophysiology of Amyotrophic Lateral Sclerosis (ALS) and Frontotemporal Dementia (FTD), in addition to Limbic-predominant age-related TDP-43 encephalopathy (LATE) Alzheimer’s disease (AD)^3-6^. The likely underlying etiology encompasses homeostatic imbalance between nuclear and cytoplasmic localization^7^, formation of aberrant inclusions composed of ubiquitinated and hyper-phosphorylated TDP-43^7-9^, and an increase in proteolytic cleavage of cytoplasmic TDP-43^10^. TDP-43 consists of an N-terminal domain (NTD), two tandem RNA recognition motifs, RRM1 and RRM2, and an intrinsically disordered prion-like C-terminal domain (CTD)^9^. It was demonstrated that the NTD and RRM domains contribute to TDP-43 pathology and possibly amyloid formation^11-13^, highlighting the need to fully understand the structural role these domains play in TDP-43 pathology.

Our laboratory, driven by the potential of TDP-43 as a target for treating neurodegenerative diseases, has focused on identifying druggable sites on TDP-43, particularly those involved in RNA/protein interactions. We identified compounds targeting the RNA-binding domain yielding rTRD01^14^ and the N-terminal domain of TDP-43 yielding nTRD22 ^15^. In the course of developing these chemical probes, we discovered potential allosteric modulation of RNA binding. nTRD22, while specifically targeting the N-terminal domain (NTD) of TDP-43 and not binding to the RRM domains, causes changes in the NMR spectrum peaks of the RRM domains and inhibits typical RNA-binding, suggesting an allosteric modulation^15, 16^. Given these observations, we formulated the hypothesis that the NTD and RRM domains might be communicating with each other either through allostery or direct interaction, as previously proposed^17^. This was supported by a recent small-angle X-ray scattering (SAXS) study showing NTD interacting with the RRM domains, although the TDP-43 construct in this study had all tryptophan residues mutated to alanine residues^18^.

In this study, we proposed the hypothesis that the N-terminal domain might be interacting closely with the RRM domain, and this interaction could play a crucial role in RNA binding, especially in the case of weaker, non-specific RNA binding. We (i) reveal, via NMR, the shift of residues in the RRM domain when the NTD is present, suggesting that the NTD is interacting with the RRM domain, (ii) employ molecular modeling to validate this as a plausible conformation, and (iii) demonstrate that RNA exhibits different binding modes depending on whether the NTD is present or absent.

## Methods

### Materials

All reagents were purchased from Sigma (St. Louis, MO, USA), Fisher Scientific (Hampton, NH) unless otherwise indicated, and IDT DNA.

### Purification of ^15^N-labeled recombinant TDP43 subdomains

Human TDP43_102-260_ and TDP43_1–260_ were expressed in *E. coli* BL21(DE3) cells (Novagen) in LB rich or M9 minimal media supplemented with ^15^NH_4_Cl. After the OD_600_ reached 0.8, 0.5 mM isopropyl β-d-1-galactopyranoside was used to induce protein expression at 20 °C for 24 h. Cells were collected by 4500 rpm centrifugation and resuspended in 40 mM Hepes, pH 7.5, 500 mM NaCl, 5 mM DTT, 30 mM imidazole, and EDTA-free protease inhibitor cocktail. Cell disruption was performed by three rounds of high-pressure homogenization at 15,000 PSI with an LM10 Microfluidizer (Microfluidics, Westwood) and cell debris was removed by centrifugation at 50,000 rpm for 1 hour at 4°C. The supernatant was then loaded on a His-Trap (GE healthcare, Uppsala, Sweden) previously equilibrated using the Lysis buffer. Protein was then eluted with a linear gradient of imidazole up to 400 mM. Eluted fractions of pure protein were dialyzed into NMR buffer (40 mM HEPES, pH 6.5, 300 mM NaCl, 4 mM DTT). Dialyzed protein was concentrated with Amicon Ultra 15 centrifugal filters (Regenerated cellulose 10,000 NMWL; Merck Millipore, Darmstadt, Germany). Protein concentration was determined by Pierce660 assay using bovine serum albumin as the standard. The purity of the eluted protein was verified by SDS–PAGE.

### Purification of recombinant TDP-43 N-terminal domain (NTD)

Human TDP43_1–102_ was expressed in *E. coli* BL21(DE3) cells (Novagen) in LB rich or M9 minimal media supplemented with ^15^NH_4_Cl. After the OD^600^ reached 0.8, 1 mM isopropyl β-D-1-galactopyranoside was used to induce protein expression at 30°C overnight. Purification was performed identically as the other TDP-43 subdomain constructs and stored in an NMR buffer.

### 2D ^15^N-HSQC-NMR

All NMR data were collected in NMR Buffer: 40 mM HEPES pH 6.5, 300 mM NaCl, and 4 mM DTT on a Bruker Avance NEO 800 Mhz spectrometer with TCI-H&F/C/N probe at 25ºC. A transverse relaxation optimized spectroscopy (TROSY)^19^ with a solvent suppression pulse sequence was used to acquire HSQC data for all ^15^N labeled His-TDP43 at 25 µM. Under the same conditions, (UG)_4_ was titrated at a 1:2 molar ratio for all constructs. NMR data processing and analysis were performed using programs NMRPipe^20^ and Sparky^21^. Chemical shift differences were calculated using the following equation^22^:

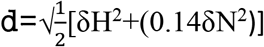

Chemical shifts that were greater than 3 standard deviations (σ) were considered significant.

### De Novo Sequence to Structure Prediction using I-TASSER-MTD and RaptorX

The partial sequences for TDP-43 (ID: Q13148) were retrieved from the NCBI protein database (https://www.ncbi.nlm.nih.gov/protein). The structure of the protein was predicted using the I-TASSER-MTD or RaptorX server. The structures were analyzed and validated by Prosa^23^ and SAVES^24^ servers. Predicted models were visualized using Pymol and ChimeraX^25^.

### In silico docking using ClusPro

The ClusPro server was used for protein-protein docking^26-28^. Briefly, ClusPro performs exhaustive sampling of ligand positions on the target and then scores by clustering the lowest-energy structures using four different scoring functions: (1) balanced, (2) electrostatic-favored, (3) hydrophobic-favored, and (4) van der Waals plus electrostatic favored. The ligand was the first NMR model of the N-terminal domain (2n4p^29^), prepared by removing the 6-His tag. The target was the first model of the RRM1-RRM2-RNA NMR structure (PDB ID 4bs2)^30^, with the 6-His tag, the C-terminal 10 amino acid linker, and the RNA molecule clipped to (UG)_4_ before docking. Default parameters were used. Docking was repeated using attractors on RRM2 based on CSP shifts, but the results were nearly identical to the runs without attractors and are thus not shown. However, obtaining similar models using different parameters lends confidence to the output^27^.

### Microscale Thermophoresis

The UG repeats (UG)_4_: rUGrUGrUGrUG and (UG)_6_: rUGrUGrUGrUGrUGrUG were purchased from IDT DNA. The UG repeats ((UG)n) were resuspended in PBS-0.05%T buffer provided by Monolith His-Tag Labeling Kit RED-tris-NTA 2^nd^ Generation (Nanotemper, Germany) to stock concentrations of 5 mM and stored at -20 °C.

HisTDP-43_1-260_, HisTDP-43_102-260,_ and the UG repeats were diluted to their working concentrations in the assay buffer (Monolith’s PBS-0.05%T). The protein was labeled according to the instruction manual provided by the manufacturer (Nanotemper, Germany). Briefly, 200 nM of the protein was labeled with 100 nM NT-647 dye of Monolith His-Tag Labeling Kit RED-tris-NTA 2^nd^ Generation (Nanotemper, Germany) and incubated at room temperature for 30 minutes followed by centrifugation at 15000Xg for 10 min at 4°C. A 16-point serial dilution of each UG repeat was prepared for the assay. 10 µL each of labeled HisTDP43 and (UG)n were mixed in a tube at different concentrations of ((UG)n). The final concentration of the labeled HisTDP-43 in the assay was 50 nM. For assays with labeled HisTDP-43_1-260_, thermophoresis signals were captured using MST premium capillaries at 60% excitation power and 40% MST power on a Monolith NT.115 device (Nanotemper, Germany). The data were analyzed with MO Affinity Analysis software (Nanotemper), utilizing the Hill model, and fitted using the specific binding with the Hill slope model in GraphPad Prism 9.1.2. For assays with labeled HisTDP-43_102-260,_ the thermophoresis signals were captured at 650 nm and 670 nm using MST premium capillaries at 80% excitation power and medium MST power on a Monolith X device (Nanotemper, Germany). The data, analyzed as a ratio of 670 nm/650 nm, were fitted using the Kd model for (UG)_6_ and the Non-Binder model for (UG)_4_ in MO Control 2 software and then plotted in GraphPad Prism 10.2.3.

## Results

### 2D [1H,15N] HSQC-NMR of TDP-43_102-260_ vs. TDP-43_1-260_

We examined the effect of NTD on RRM domains by comparing the 2D [^1^H,^15^N] HSQC-NMR spectra of two TDP-43 constructs, one containing the RRM domain with NTD and one without the NTD; **(Figure 1**). For the remainder of the manuscript we refer to these constructs as follows: TDP-43 that covers only the RRM domains (RRM1 and RRM2) from 102-260 amino acids is designated TDP-43_102-260_; the construct that covers the N-terminal domain and the RRM domain will be TDP-43_1-260_ and covers amino acids 1-260 and the construct that covers only the N-terminal domain 1-102 as TDP-43_1-102_ (**Supplemental Figures 1, 2**). TDP-43_102-260_ spectra reproducibly display a well-folded protein indicated by the sharp, well-resolved peaks as seen before with 143 identified resonances^31^ (**Figure 1, Supplemental Figure 3A)**. TDP-43_1-260_ contains three folded domains that include the NTD, RRM1 and RRM2-based on this, we would expect ∼222 amide peaks in the ^15^N-HSQC-NMR acquired spectra. For the first time, we show a high-resolution 2D [^1^H,^15^N] TROSY-HSQC spectrum of multiple TDP-43 domains connected in tandem by acquiring TDP-43_1-260_ in low protein concentrations resulting in 208 combined resonances from the NTD and RRMs **(Supplemental Figure 3A-C)**. Superimposing TDP-43_102-260_ NMR spectrum onto TDP-43_1-260_ results in overlapping peaks from the RRM region, but also distinguishable peaks from NTD resonances **(Figure 1A)**. By transferring ^1^H,^15^N assigned resonances of apoTDP-43 RRM domains to the superimposed spectra, we found significant chemical shifts primarily of charged residues in the RRM domains **(Figure 1B)**. Chemical Shift Perturbation (CSP) analysis shows significant clustered shifts in the ribonucleoprotein sequences of RRM1 and RRM2 **(Figure 1C)**. Next, we mapped the chemical shifts onto the RNA-bound tandem RRM structure revealing key residues involved in RNA-binding that are appreciably perturbed by the presence of the NTD in TDP-43_1-260_ **(Figure 1D)**. Of importance to ALS and mutations found in TDP-43 connected to ALS, one of the most significantly shifted residues is D169, which is a mutated to a Glycine residue in ALS patients (**Figure 1B, D**), previously reported to (i) disrupt the ATP-binding capacity of the RRM1 domain^32^, (ii) disrupt phase separation-induced by NEAT1 RNA-binding to TDP-43^33^, and (iii) was shown to increase thermal stability of a TDP-43(1-265) construct^34^.

**Figure 1.**
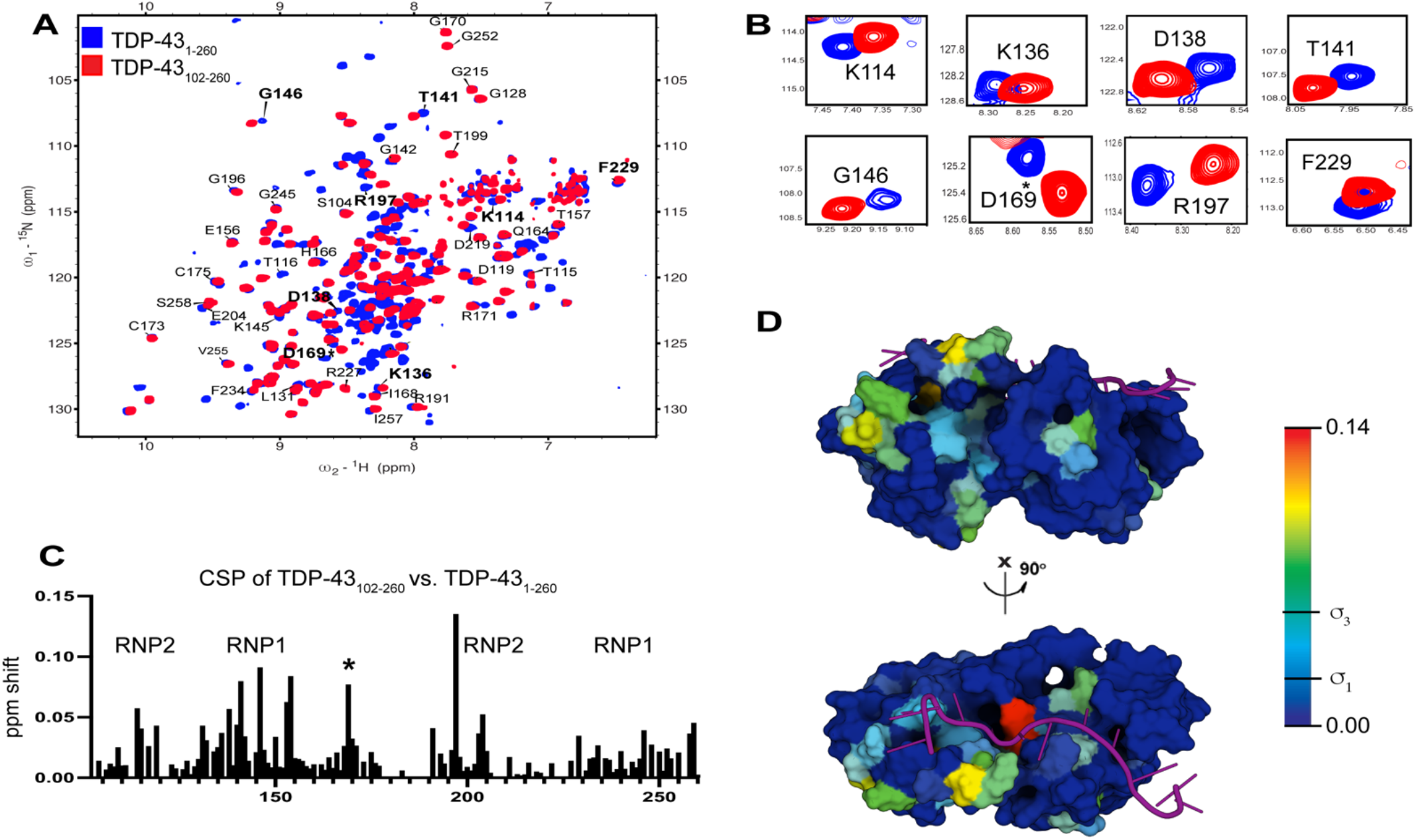
2D HSQC-NMR of TDP-43_1-260_ identifies NTD-RRM interactions. **(A)** 2D ^15^N-HSQC-NMR overlay of ^15^N and ^1^H resonances of TDP-43_102-260_ (red) onto ^15^N TDP-43_1-260_ (blue) at 25 µM under identical conditions. **(B)** Zoomed regions of 2D HSQC-NMR overlay show exemplar polar residues of the RRM domains that are shifted in the presence of the NTD. **(C)** Chemical Shift Plot (CSP) from resonances of TDP-43_102-260_ vs TDP-43_1-260_ shows shifts in the RNP-1 and RNP-2. RNP-1 and RNP-2 are short motifs in the RRM domains known as ribonucleoprotein domains ^1^.**(D)** CSP analysis was projected as a heat map representative of shift intensities onto the tandem RRM RNA-bound structure (PDB 4BS2)^2^.

### Prediction of interdomain contacts by simulating protein-protein interactions

To further understand how the NTD interacts with RRM domains, structures of the NTD (PDB: 5MRG^35^) and RRMs (PDB: 4BS2^30^) were simulated as protein-protein interactions using High Ambiguity Drive protein-protein Docking (HADDOCK)^36, 37^. HADDOCK molecular modeling of protein-protein interactions is unique in that it uses Ambiguous Interactions Restraints (AIRs) from experimental data in combination with three-step simulated interaction using rigid-body docking, semiflexible refinement, and minimized by molecular dynamics simulations in an explicit solvent shell. Using default settings, TDP-43 structures along with predefined “active” residues from the CSP analysis were submitted to the HADDOCKv2.4 web server. HADDOCK assembled 92 structures into 13 clusters, representing 46% of all water-refined models generated. The results were ranked using the standard energy scoring metric in HADDOCK and three clusters, #2, #3, and #5 containing 25 structures showed favorable HADDOCK scores with negative Z-Score values **(Table 1)**.

**Table 1.** Results of top three clusters from HADDOCK protein-protein docking simulations of TDP-43_1-102_ and TDP-43_102-260_.

| HADDOCK Results | HADDOCK Cluster 2 | HADDOCK Cluster 3 | HADDOCK Cluster 5 |
| --- | --- | --- | --- |
| HADDOCK score | -83.5 +/- 3.3 | -82.0 +/- 11.2 | -75.7 +/- 7.7 |
| Cluster size | 9 | 8 | 7 |
| RMSD from the overall lowest-energy structure | 7.3 +/- 0.1 | 0.9 +/- 0.5 | 5.4 +/- 0.3 |
| Van der Waals energy | -61.0 +/- 7.6 | -66.2 +/- 1.2 | -54.5 +/- 6.1 |
| Electrostatic energy | -221.3 +/- 22.8 | -262.1 +/- 63.5 | -242.7 +/- 45.7 |
| Desolvation energy | 16.7 +/- 1.4 | 27.2 +/- 2.9 | 24.2 +/- 2.3 |
| Restraints violation energy | 51.1 +/- 37.9 | 93.9 +/- 35.6 | 30.5 +/- 30.2 |
| Buried Surface Area | 1691.5 +/- 31.2 | 1921.3 +/- 34.4 | 1585.3 +/- 125.2 |
| Z-Score | -1.6 | -1.4 | -0.8 |

Visualization of these docking results predicts NTD interacting with both RRM domains, but with a tendency towards RRM1 **(Figure 2A)**. A notable pattern emerges when comparing clusters of HADDOCK Score against RMSD and total electrostatic energy, where the most favorable clusters exhibit the lowest electrostatic potential energy. This underscores the significance of electrostatic residues, as observed in the 2D ^15^N-HSQC-NMR, which become apparent with the addition of the NTD to the construct^38^ **(Figure 2B)**. Of these top clusters, #2 and #5 were analogous in values for HADDOCK Score, RMSD, and electrostatic energy with both results stacking the NTD onto RRM1, while cluster #3 shows the NTD stacking onto RRM2 **(Figure 2C, D)**. From our CSP analysis, K136, D138, T141, and G146 of RRM1 along with R197 and F229 of RRM2 were the most significantly perturbed residues and most likely sites of NTD interactions. Using the heat map projected onto the RRM structure from our CSP analysis, visualization of all HADDOCK clusters correlated well with the predicted interactions of both RRM domains **(Figure 2E, F)**. Using these same structures, we repeated the NTD-RRM interactions using the ClusPro protein-protein webserver^26^. ClusPro only uses rigid-body docking with inter-and intramolecular interaction energies evaluated by force field calculations. ClusPro results also indicated NTD-RRM interdomain contacts, but all force field weights including electrostatic, hydrophobic, or balanced replicated NTD stacking onto RRM2 **(Supplemental Figure 4)**^27, 39^.

**Figure 2.**
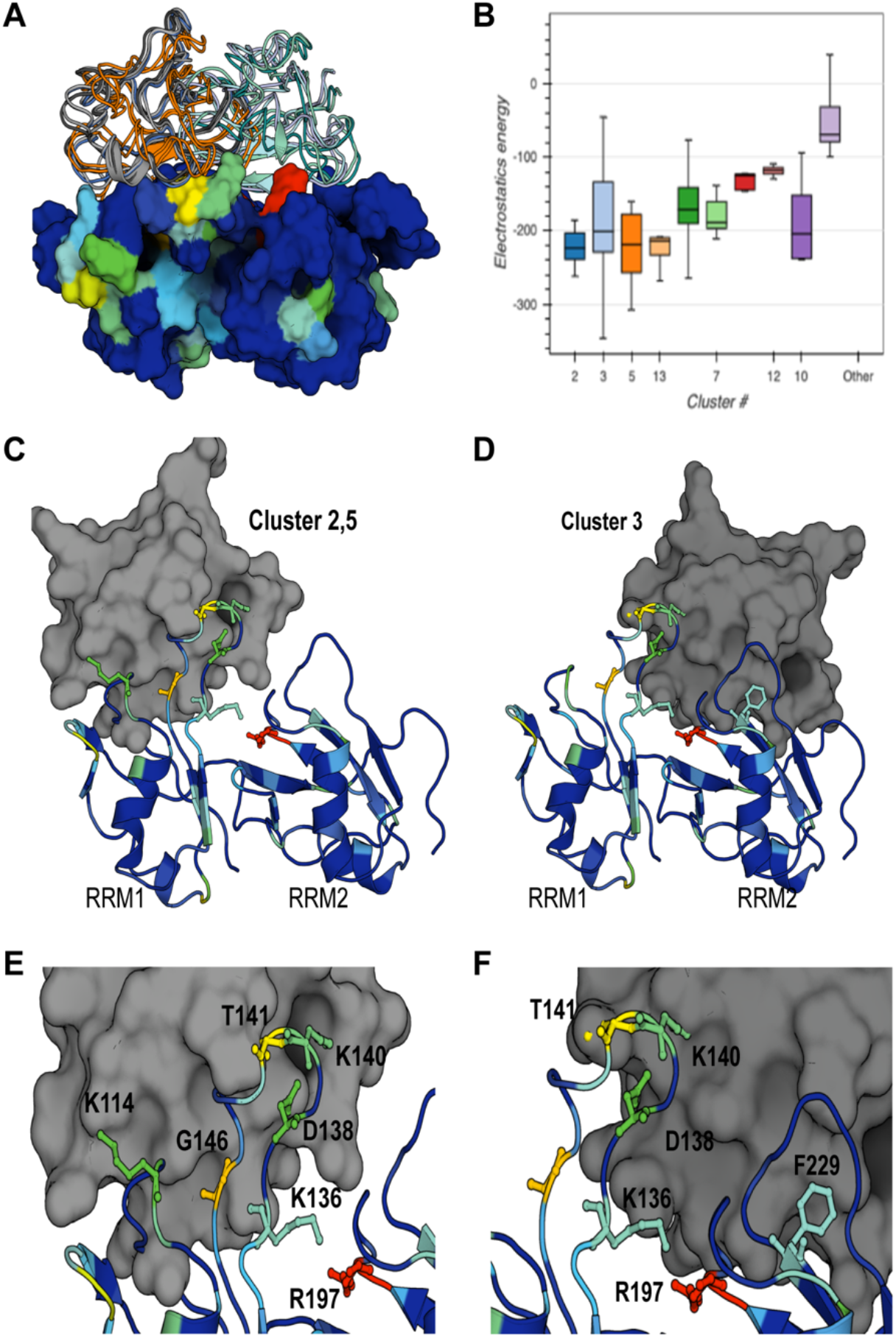
HADDOCK protein-protein simulations of NTD-RRM stacking. **(A)** Top 3 HADDOCK clusters contain 25 structures stacking the NTD (cartoons) onto the RRM domains (surface). The CSP heat map from 2D HSQC-NMR is used to show correlations between experimental and simulated data. **(B)** HADDOCK results were ranked by HADDOCK Score, Z-Score, RMSD, and electrostatic potential energy. A general trend was seen that the lower-RMSD containing top clusters also had the lowest potential electrostatic energy. **(C)** HADDOCK Clusters 2 and 5 stacks the NTD (grey) onto RRM1 **(D)** Cluster 3 stacks NTD (grey) onto the RRM2 domain **(E)** Zoom of Clusters 2 and 5 stacked onto RRM1 with significant shifts from CSP analysis labeled. **(F)** Zoom of Cluster 3 stacked onto RRM2 with significant shifts from CSP analysis labeled.

### Biophysical assessment of NTD-RRM interactions using CPMG-NMR and SPR

The transient formation of protein-protein interactions is challenging to biophysically characterize due to their dynamic nature and shallow interfaces. This complexity is further amplified by the differential behavior of TDP-43 constructs across various pHs (**Supplemental Figure 2**). NMR relaxation experiments excel in this area, as they can report time scales ranging from picoseconds to milliseconds^40^. As domain motion of large multidomain proteins occurs in the µs-ms timescale, Carr-Purcell-Meiboom-Gill (CPMG)– relaxation dispersion NMR spectroscopy is a useful approach for measuring protein dynamics over a range of timescales^41, 42^. CPMG-NMR quantifies the line broadening that occurs by the chemical exchange between the ground state and one or more excited states of NMR resonances over specific time points^43, 44^. 2D ^15^N-HSQC-NMR spectra of 100µM TDP-43_102-260_ were acquired with a CPMG-filter of 20 ms, 40 ms, 70 ms, 140 ms, 270 ms, and 540 ms. Among these, the 20 ms, 70 ms, and 140 ms displayed significant changes in signal intensity and are appropriate ranges for relaxation rates of 20kDa proteins^41^. Using these same parameters, unlabeled NTD_1-102_ was added to ^15^N TDP-43_102-260_ (RRM1 and RRM2) at a 1:4 (NTD: RRM1 and RRM2) molar ratio and showed significant changes in signal intensity compared to apoTDP-43_102-260_ **(Figure 3A)**. As the NTD readily oligomerizes, concentrations above a 1:4 molar ratio of NTD:RRM12 showed no signs of interaction and were likely necessary to compete the oligomerization of NTD. Integration of each peak provides a local relaxation constant and averaging all relaxation constants of a given spectrum provides the global tumbling rate of proteins in solution (T2). In doing so, we observed evidence of complex formation by a measured increase of the T2 tumbling rate of TDP-43_102-260_ in solution with the NTD_1-102_ **(Figure 3B, C)**.

**Figure 3.**
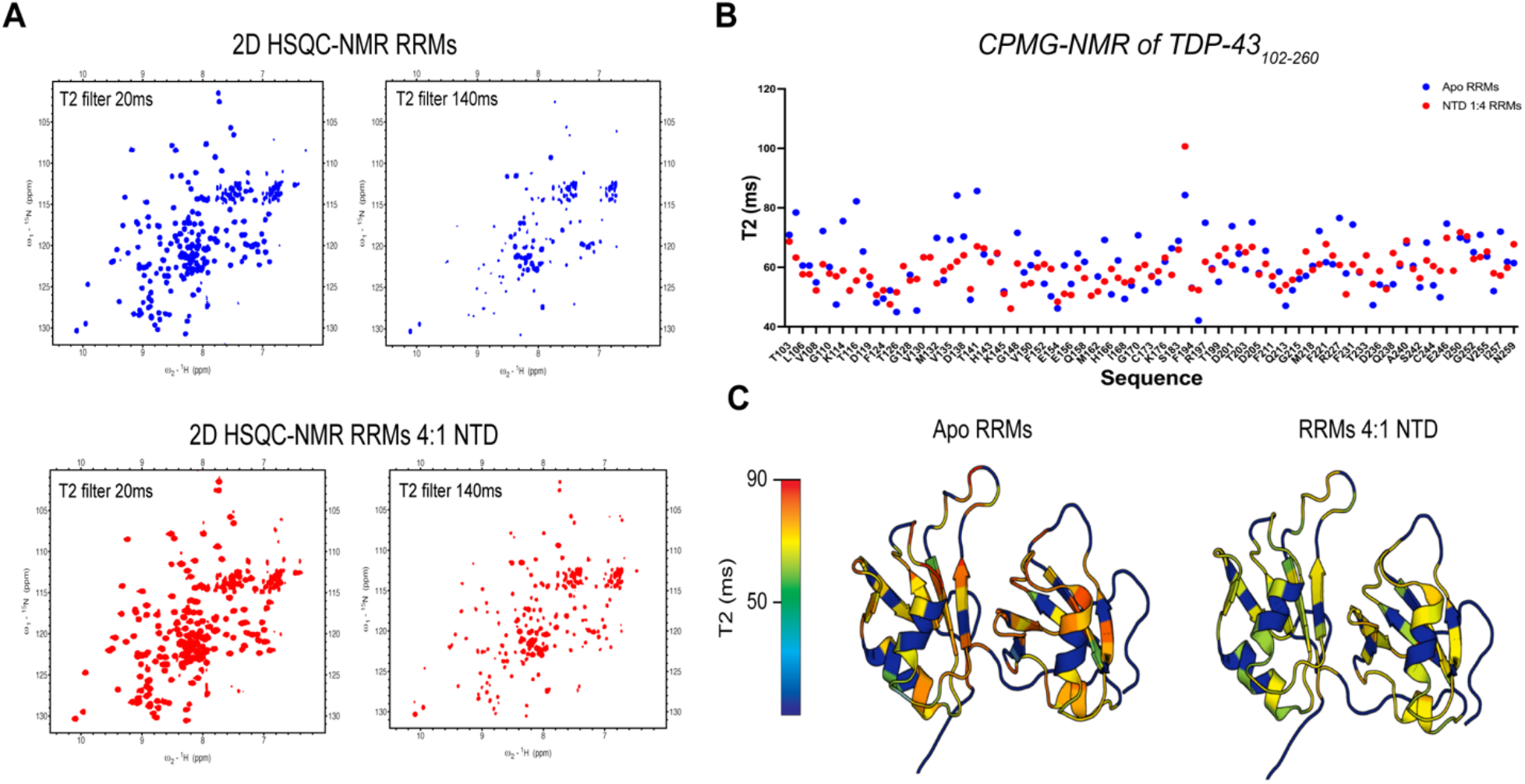
CPMG-NMR and SPR of protein-protein interactions between TDP-43_102-260_ and TDP-43_1-102_. **(A)** CPMG-NMR was acquired from ^15^N-labeled TDP-43_102-260_ at 100 µM with five T2 filters ranging from 20-540ms. 20ms and 140ms spectra are shown as representative of the decay rate for apoRRMs. **(B)** 25 µM unlabeled NTD_1-102_ was added to ^15^N TDP-43_102-260_ and the same T2 filters were acquired using CPMG-NMR with five-time points. 20ms and 140ms spectra are shown as representative of the decay rate for apoRRMs. **(C)** Global T2-relaxation constants were calculated for apo and bound TDP-43_102-260_ at each T2 filter, showing a significant increase in the tumbling rate.

### NTD competes with RNA-binding to the RRM domains in 2D ^15^N-HSQC-NMR

Given that the NTD_1-102_ showed interaction with the RRM domains and was proposed to engage with crucial residues involved in RNA binding, we further explored the potential relationship between the interaction of NTD-RRMs and RNA-binding. We identified specific differences between short RNA sequences binding to TDP-43_102-260_ of the RRMs alone vs TDP-43_1-260_ that included the NTD. Titration of a short 5’-UGUGUGUG-3’ (henceforth referred to as: (UG)_4_) RNA onto TDP-43_102-260_ and observation of changes using 2D ^15^N-HSQC-NMR resulted in no chemical shifts **(Figure 4A)**. (UG)_4_ has two fewer nucleotides (eight) than that of the total number of RNA interaction sites of RRM1-RRM2 (ten) and in our experimental design was not found to bind to both RRM domains in tandem. Given that TDP-43 RRM domains exhibit a greater affinity for extended UG-rich RNA repeats, which also promote cooperative RRM-RNA binding, we propose using a shorter UG-motif. This motif would fall within the suitable affinity range for 2D _15_N-HSQC-NMR observation and could enhance NMR resolution by reducing the number of interactions among multiple TDP-43 units. Another possibility is that the 2D ^15^N-HSQC-NMR results show a slow exchange rate between (UG)_4_ and TDP-43_102-260_, as is normally seen with low nM affinity ligands that generally go undetected by chemical shift analysis^40, 45-47^.

**Figure 4.**
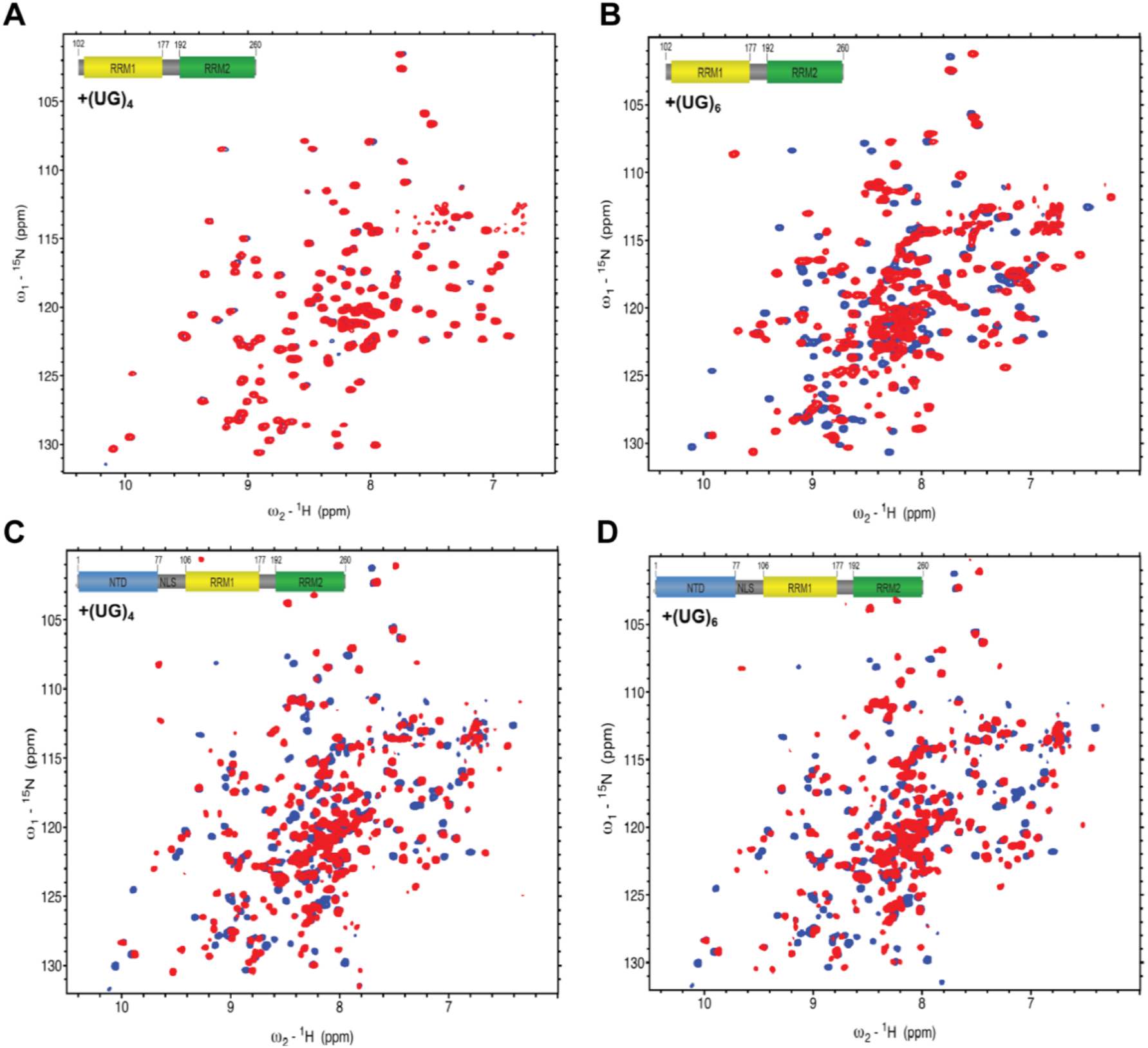
2D ^15^N-HSQC-NMR identifies a change in RRM-RNA binding dynamics in the presence of the N-terminal Domain. **(A)** 2D ^15^N-HSQC-NMR overlay of apoTDP-43_102-260_ (blue) and (UG)_6_-bound (red) at 1:1.**(B)** 2D ^15^N-HSQC-NMR overlay of apoTDP-43_102-260_ and (UG)_4_-bound (red) at 1:1. **(C)** 2D ^15^N-HSQC-NMR overlay of apoTDP-43_1-260_ (blue) and (UG)_6_-bound (red) at 1:1. **(D)** 2D ^15^N-HSQC-NMR overlay of apoTDP-43_1-260_ and (UG)_4_-bound (red) at 1:1. Significant chemical shifts are observed in RRM residues and NTD residues, indicating NTD plays a role in RNA binding and a significant change in the exchange rate between RNA-TDP-43 binding.

We tested the binding of RNA of different length to several TDP-43 protein constructs using Microscale Thermophoresis (MST). MST allows the detection of interaction between a fluorescently labeled target (His-TDP-43 here) and a ligand ((UG)n) under an infrared laser that generates a temperature gradient in capillaries containing samples. During a binding event, alterations in the hydration shell of biomolecules lead to a relative change in their movement along the temperature gradient generating an altered thermodiffusion and a change in the MST signal. A variation in the MST signal can provide an apparent affinity constant, Kd, by using different concentrations with a Monolith X. This device, equipped with spectral shift technology, can identify any alterations in the fluorescence emission of a fluorescently tagged target at wavelengths of 650 nm and 670 nm, which are indicative of a binding event. Fluorescence calculated at 670/650 nm and plotted against the ligand concentrations would yield an affinity constant. MST confirmed no binding of (UG)_4_ to the TDP-43_102-260_ **(Figure 5A, B)**. Strikingly, titration of the same RNA onto TDP-43_1-260_ shows a large binding affinity with an apparent Kd of 350 nM (**Figure 5C, D**), indicative of large conformational changes and consistent with the fast-exchange observed by NMR (**Figure 4C-D**).

**Figure 5.**
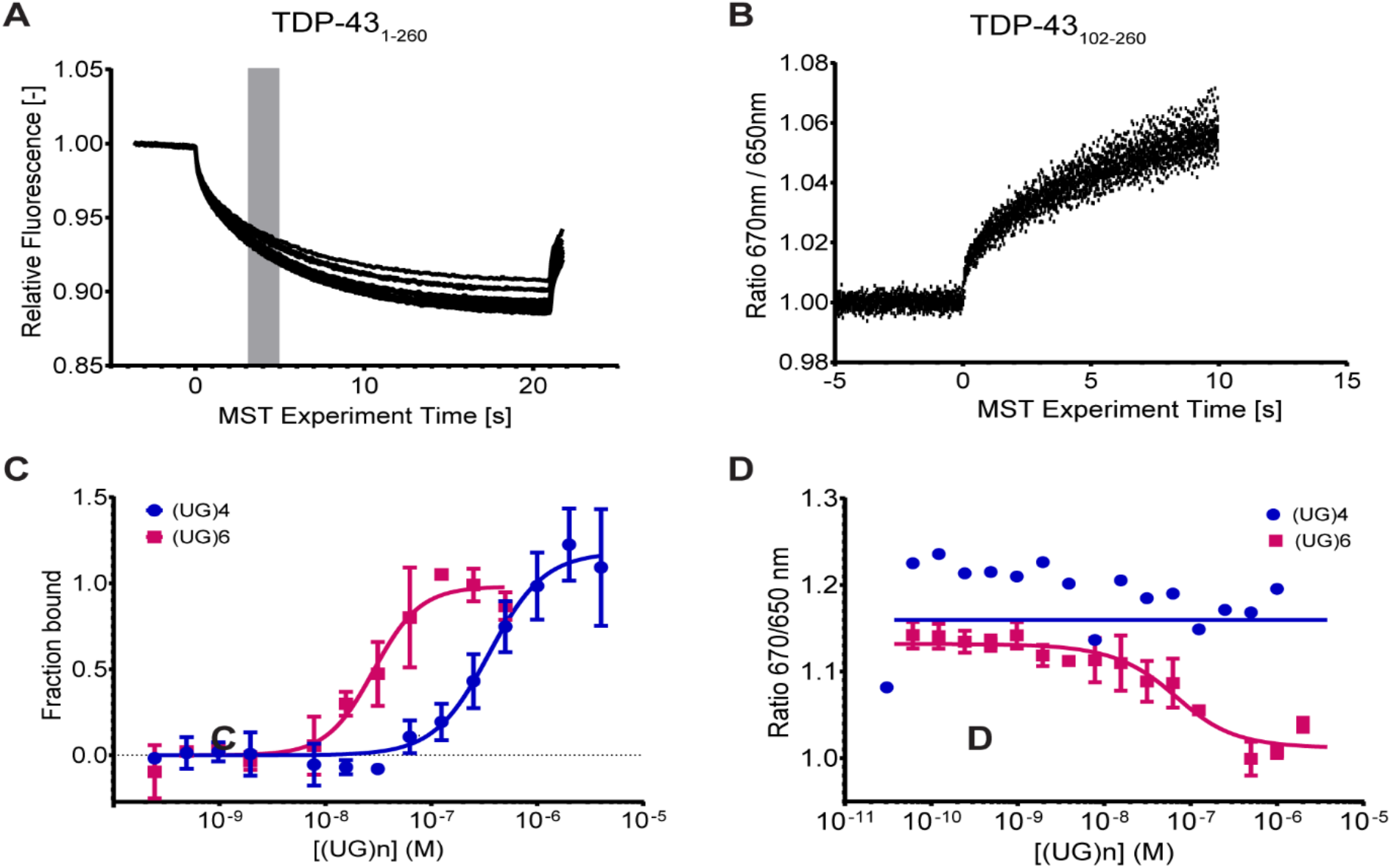
MST Analysis of RNA binding to TDP-43 constructs. (A) A representative set of thermographs of (UG)_n_ binding in increasing concentrations (0.24 nM to 4000 nM) to the NT-647 labeled HisTDP-43_1-260_ showed altered thermodiffusion, resulting in well-defined curves. The grey bar (at ∼5 s) indicates the steady-state time point used for analyzing the MST measurements in the graphs displayed in panel B. (B) MST values were used to determine the dissociation constant for binding of (UG)_n_ to HisTDP-43_1-260_. The data were fitted as described in the Methods section yielding an apparent Kd of (UG)_4_ = 246.5 nM with a confidence interval (CI) of 178.0-345.5 nM and that for (UG)_6_ = 28.42 nM with CI of 21.28-37.7 nM. (C) This representative set of thermographs shows the thermophoretic movement of (UG)6 when bound to NT-647 labeled HisTDP-43_102-260_ at increasing concentrations (0.006 nM to 2000 nM). The fluorescence signals were recorded at 650 nm and 670 nm, and their ratio plotted against the increasing concentrations of (UG)_6_ showed altered thermodiffusion, resulting in well-defined curves. (D) MST values taken at a 670 nm/650 nm ratio were used to determine the dissociation constant for (UG)_n_ binding to HisTDP-43_102-260_. The data were fitted using MO Control 2 software as described in the Methods section, resulting in an apparent Kd of 39.8 nM for (UG)6, with a CI of 16.9-93.8 nM. For (UG)_4_, the MST values did not produce a binding curve or fit the Kd model. Instead, they fit the Non-Binder model, indicating no binding interaction between HisTDP-43_102-260_ and (UG)_4_. Data is represented by Mean ± SD for n=3 except for (UG)4 in panel D where n=1 is represented.

To corroborate this further, we also tested the binding of a slightly longer UG-motif (UG)_6_ which is four nucleotides longer than (UG)_4_ and two nucleotides longer than the total number of RNA interaction sites of RRM domains as a positive control for RNA-binding event. We found that (UG)_6_ was successfully bound to both the TDP-43 constructs with a 10-fold increase in the binding affinity as compared to (UG)_4_ (**Figure 4,5**). These data suggest the following two possibilities (1) in the absence of NTD, longer RNAs bind to the RRM domains covering the RNA interaction sites, and (2) shorter RNA binds to the RNA interaction sites only in the presence of NTD.

### De novo sequence to structure prediction of TDP-43_1-260_

There have been recent advancements in the template and non-template-based *de novo* sequence to structure predictions of multidomain proteins including RosettaFold, RaptorX, and I-TASSER^48,49^. Iterative Threading ASSEmbly Refinement (I-TASSER) is a computational approach that identifies structural templates from the PDB and produces full-length atomic models by iterative template-based fragment assembly. Most recently, I-TASSER-MTD was released as a pipeline specially designed to automatically generate high-quality structures of proteins containing multiple domains from amino acid sequences alone by combining I-TASSER with inter-domain distances and contacts predicted by a deep convolutional neural network called DeepPotential. Submitting the TDP-43_1-260_ sequence through I-TASSER-MTD shows several conformations of the NTD stacking onto RRM1, concurrent with HADDOCK results (**Supplemental Figure 5A**). To build further on this, we also used RaptorX as a non-template contact-based *de novo* protein structure prediction. Intriguingly, RaptorX results converged to NTD stacking onto RRM2, more closely resembling the ClusPro results than the HADDOCK or I-TASSER-MTD results **(Supplemental Figure 5B)**. The outcomes are likely to vary between the contrasting setups. I-TASSER-MTD, a template-based method, models the tandem RRM domains in a manner closely resembling PDB 4BS2. On the other hand, RaptorX, which does not use any templates, provides greater flexibility in the connection between the RNA binding domains.

## Conclusion

The work presented here elucidates the interaction between the N-terminal domain (NTD) of TDP-43 and its RRM domains. When comparing the RRM domain in a construct where the N-terminal domain is present, the 2D ^15^N-HSQC resembles an RNA-bound form. This suggests that stacking of the N-terminal domain onto the RRM domain affects the same residues that would typically engage in RNA binding. This finding implies that the NTD may protect the RRM domain from non-specific RNA binding.

To delve deeper into the finding that suggests an interaction between the NTD and the RNA-binding residues of the RRM domains, we examined the impact of the NTD on the RRM and RNA binding through methods such as modeling, NMR spectroscopy, and MST. We acquired 2D [^1^H,^15^N] HSQC-NMR spectra of tandem RRM domains TDP-43_102-260_ and TDP-43_1-260_, a construct containing the RRMs and NTD connected by TDP-43’s nuclear localization sequence. Superimposing the HSQC-NMR spectrum of TDP-43_102-260_ onto TDP-43_1-260_ revealed significant chemical shifts within the RNA binding sites of the RRM domains. Of relevance to mutations associated with ALS is D169G, which had the greatest shift in the 2D ^15^N-HSQC when the NTD stacks on the RRM domains. This residue is significant as it is among the few identified mutations in the RRM domains linked to the pathophysiology of ALS. It has been demonstrated that this mutation enhances the stability of a construct similar to the one we utilized here (1-265 aa)^34^. Increased stability brought about by the mutation might thus be attributed to a more stable NTD stacking.

To further explore the N-terminal stacking onto the RRM domain, we modeled the NTD_1-102_ (PDB: 5MRG) and apo RRM’s amino acids 102-260 (modified PDB: 4BS2) protein-protein interactions using High Ambiguity Driven protein-protein Docking (HADDOCK) and the ClusPro protein-protein docking server. These simulations showed NTD stacking onto RRM residues with high similarity to the shifted residues observed in NMR. Additionally, de novo sequence-to-structure prediction using ITASSER-MTD and RaptorX also indicated interdomain stacking between the NTD and RRM domains. Based on the modeling result, it is not entirely clear if NTD stacks on RRM1, RRM2 or overlapping these two domains. Our CSP analysis from the NMR data comparing the NTD-containing construct to the RRM constructs seems to imply that it is skewed towards the RRM1 domain, with slight shifts of RRM2. However, the shifted residues were not assigned using multi-dimensional NMR experiments which could lead to errors of precisely which residues were most significantly shifted.

We used Carr-Purcell-Meiboom-Gill (CPMG)-NMR to verify the specific interaction between these two domains, beyond their tandem connection. We observed an increased T2 relaxation of TDP-43_102-260_ when NTD_1-102_ was present, indicating higher molecular weight and complex formation, which further supports the interaction between the NTD and the RRM domain.

Considering the substantial therapeutic potential in targeting TDP-43 and its pivotal role in the pathophysiology of numerous neurodegenerative diseases, it is crucial to understand how the apo structure of TDP-43 diverges from the RNA-bound structure and how the NTD domain can alter TDP-43 function. With the new data underscoring the significance of the NTD’s interaction with the RRM domain, we are led to question if the NTD could impact RNA binding.

We examined the potential influence of the NTD on RNA binding. We hypothesized three possible outcomes of the NTD stacking on the RRM domain: (i) it could inhibit RNA binding, (ii) it could prevent weaker RNA binding without affecting canonical UG-rich RNA binding, or (iii) it could enhance the binding of UG-rich RNA. We tested a short RNA (8 nucleotide, (UG)_4_) and found that it binds differently to the RRM domain and the NTD-RRM construct. NMR and MST data showed that (UG)_4_ does not bind to the RRM domain (TDP-43_102-260_), but it does bind to the NTD-RRM construct (TDP-43_1-260_). In contrast, (UG)_6_ binds to both constructs. This suggests that the NTD does not inhibit RNA binding, as (UG)6 binds to NTD-RRM in the same way it does to the RRM domain. Interestingly, the (UG)4 RNA, which should bind to the RRM domain, does not. However, it does bind to the NTD-RRM domain. Based on our current data, we cannot determine whether the NTD enhances the binding of the (UG)_4_ RNA or if the increased affinity is due to residues in the loop region that connect the NTD to the RRM domain (residues 77-102), which are heavily charged. This implies that the RRM alone has sufficient affinity to bind to short RNAs, with an increase in charge either from the loop region on the N-terminal domain or possibly from charged residues on the C-terminal region of TDP-43 not included in our studies (9 additional amino acids are included in the original structures of TDP-43 102-269 with RNA^30^).

Our study highlights how opening the TDP-43 NTD exposes the RRM domain and highly charged residues, particularly in the loop connecting the NTD and RRM, that then allow RNAs to bind with higher affinity and specificity. We show that there are different outcomes to binding small RNAs when the N-terminal domain stacks on the RRM. This may be important to regulate or prevent non-specific interactions of TDP-43 with RNAs of low affinity and length. This implicates a significant impact on studies that use tagged N-terminal domain constructs that may disrupt this protective stacking and shift the protein towards a more pathological form of TDP-43. In turn, this may drive aggregation otherwise prevented by constitutive N-terminal domain stacking. Indeed, one interpretation of overexpressed tagged TDP-43 and its predilection for aggregation could be due to non-specific RNA and possibly charged protein binding in the absence of N-terminal domain stacking. Future studies should thus focus on understanding if and how the N-terminal domain protects against aggregation while moving forward with a novel caveat for studies that use N-terminal tagged TDP-43.

## Supplemental Figures

**Supplemental Figure 1.**
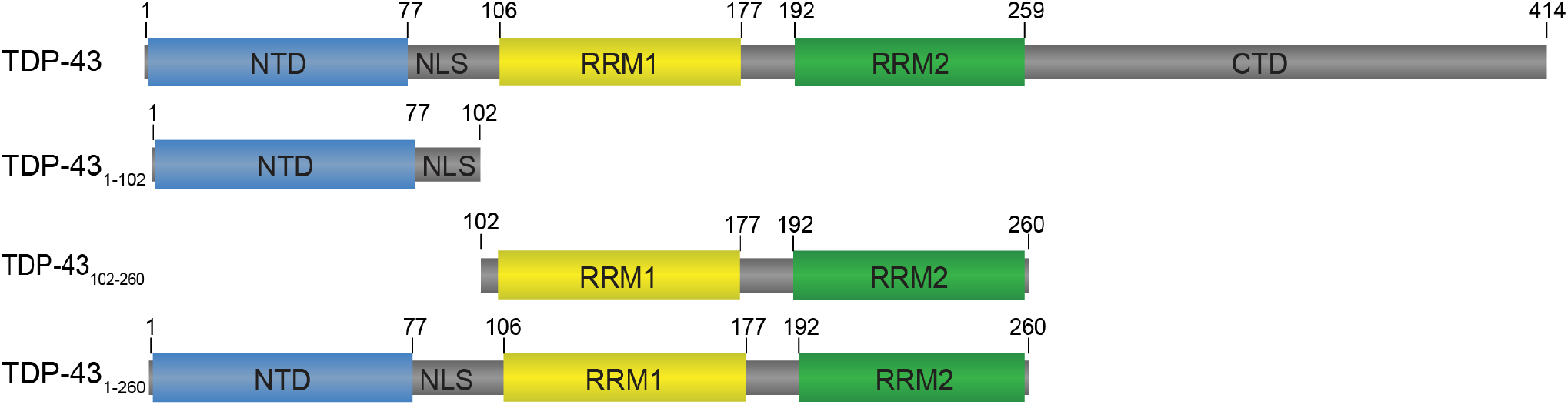
Domain architecture of subdomain constructs of TDP-43.

**Supplemental Figure 2:**
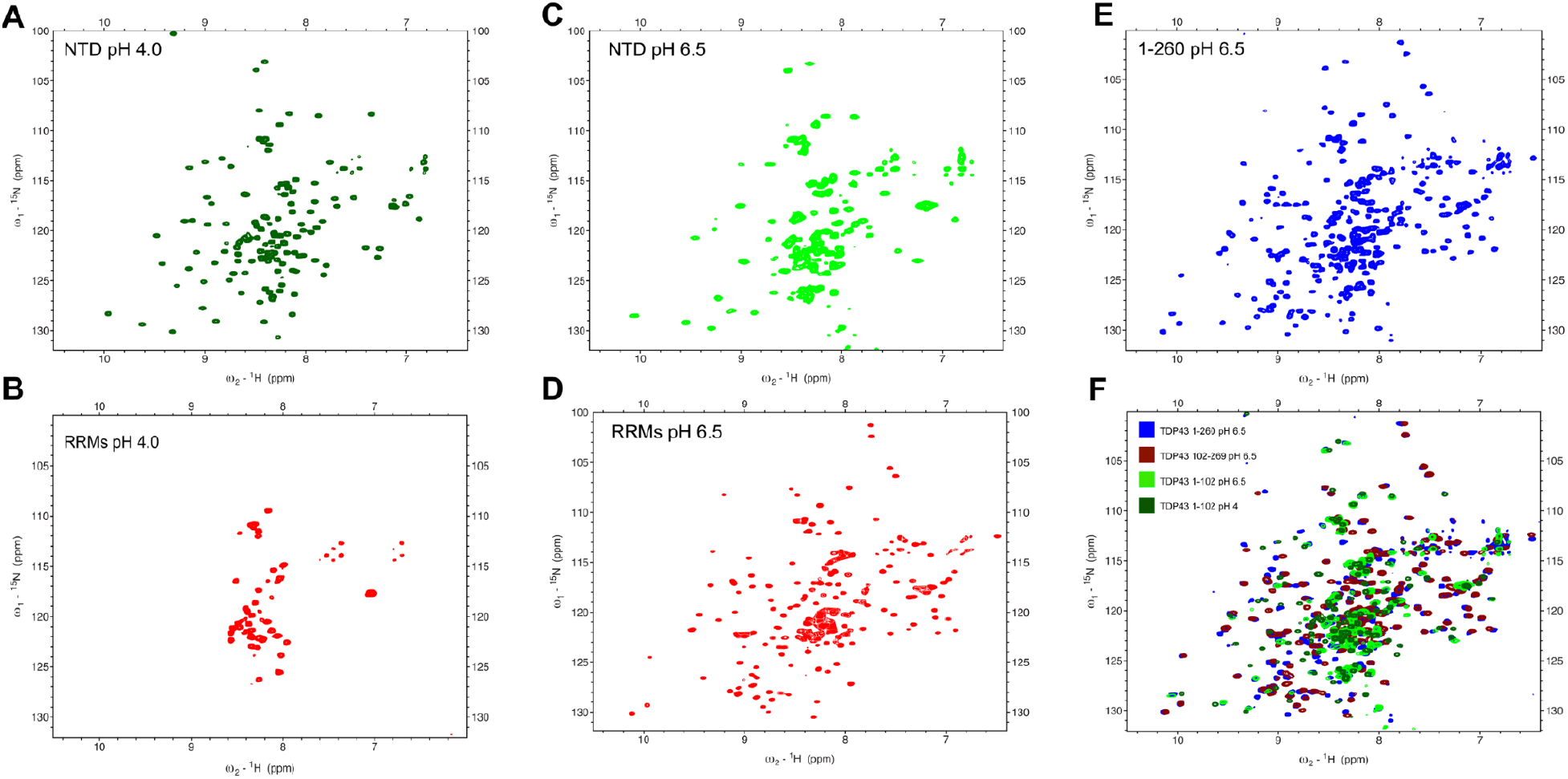
pH stability of TDP-43s NTD_1-102_, RRMs_102-260_, and a multidomain construct TDP-43_1-260_. The pH of different constructs were tested. **A**. NTD (1-102aa) at pH 4.0 **B**. The RRM domain was tested at pH 4.0 (102-260 aa) **C**. NTD (1-102 aa) at pH 6.5 D. The RRM domain (1-260 aa) at pH 6.5 **E**. NTD-RRM (1-260 aa) domain at pH 6.5 and **F**. Overlay of the NTD-RRM (1-260 aa) at pH 6.5(blue) with TDP-43 RRM (102-269) at pH 6.5(burgundy) with TDP-43 NTD (1-102 aa) at pH 6.5 (green) with the NTD (1-102) at pH 4.0 (dark green).

**Supplemental Figure 3.**
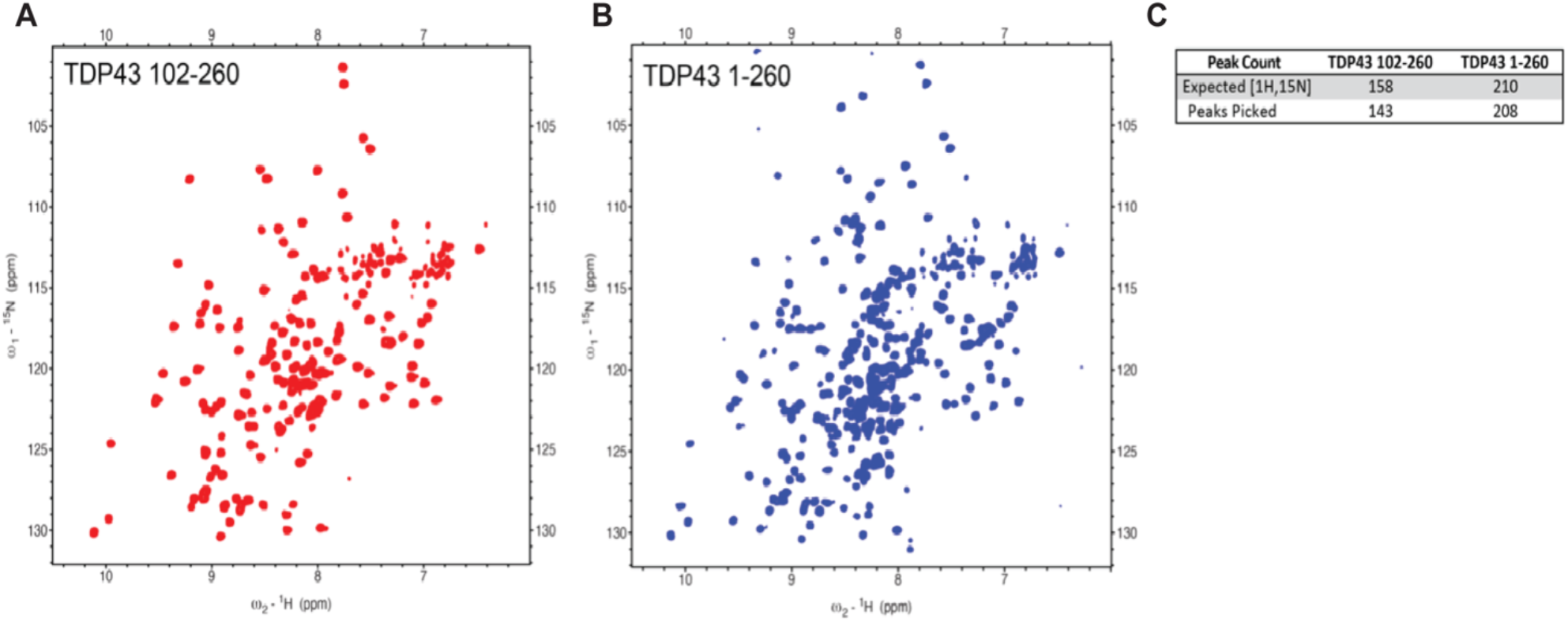
2D [1H,15N] TROSY HSQC-NMR of TDP-43 subdomains. (A) TDP-43_102-260_ acquired at 25μM in NMR buffer shows a highly resolved spectrum with well-dispersed sharp peaks. (B) TDP-43_1-260_ shows high resolution at 25μM resulting in well dispersed and sharp peaks for both the RRM domains and NTD **(C)** Expected peak count of TDP-43 sequences is calculated from sequence length subtracted by the total number of proteins and linkers. Manual peak picking excluded side-chain resonances and identified ∼95% of expected peaks for TDP-43.

**Supplemental Figure 4.**
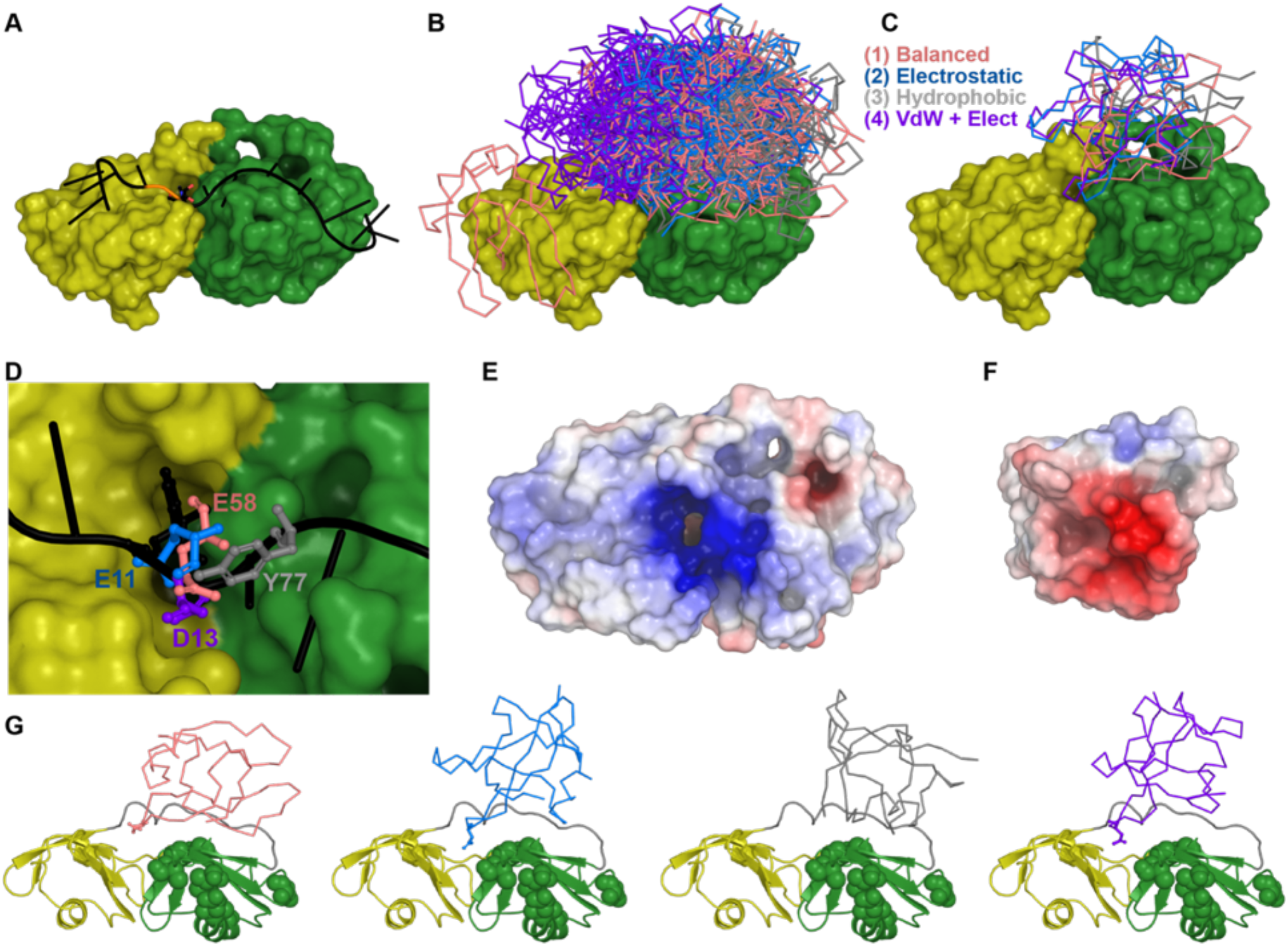
ClusPro docking of NTD to RRM1-RRM2. **A**. NMR structure of RRM1 (yellow) and RRM2 (green) shown in surface representation with RNA as a cartoon (PDB ID 4bs2 _2_). **B**. All models of docked NTD (PDB ID 2n4p ^29^) are shown as ribbons, colored by scoring function: (1) balanced models in pink, (2) electrostatic-favored models in blue, (3) hydrophobic-favored models in gray, and (4) van der Waals (VdW) + electrostatic favored models in purple. **C**. Top models for all four scoring methods. **D**. Location of residues overlapping with the RNA C’5 site. **E**. Electrostatic surface of RRM1-RRM2 and **F**. of NTD, where NTD was rotated 180 degrees about the vertical to show the surface that contacts RRM1-RRM2. **G**. Comparison of top model positions to chemical shift perturbations (green spheres).

**Supplemental Figure 5.**
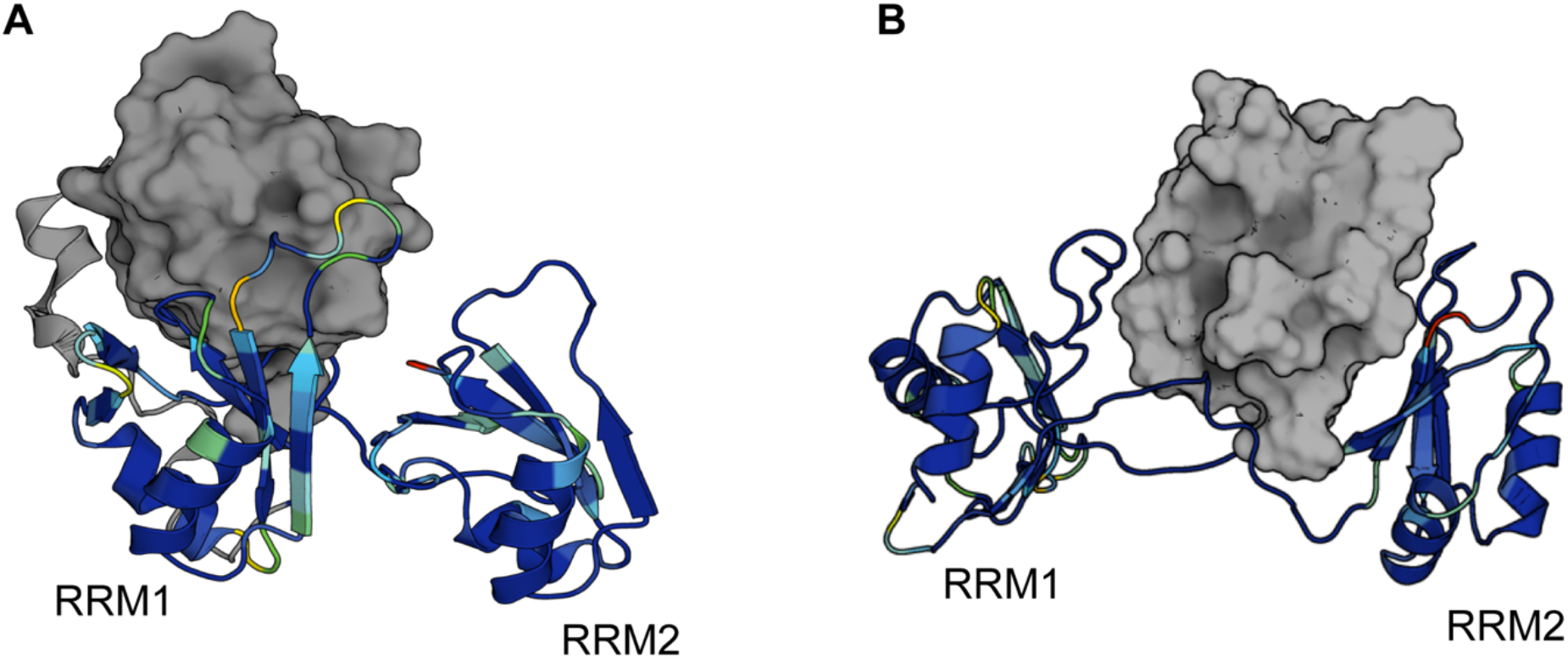
I-TASSER and RaptorX *de novo* sequence to structure prediction converges to NTD-RRM interdomain stacking. (A) TDP-43_1-260_ sequences were submitted to the I-TASSER-MTD web server. Results predicted folded NTD and RRM domains with template-based predicted interdomain stacking between the NTD and RRM1 domain. (B) RaptorX non-template contact-based structure prediction resulted in interdomain contacts between NTD and RRM2. These similar results embody the dynamic interactions between this multidomain construct. *A heat map is projected onto the predicted residues 102-260 of each model to exemplify experimental NMR data*.

